# Discovery of *NANOG* enhancers and their essential roles in self-renewal and differentiation in human embryonic stem cells

**DOI:** 10.1101/2024.12.21.628413

**Authors:** Jielin Yan, Renhe Luo, Bess P. Rosen, Dingyu Liu, Wilfred Wong, Christina S. Leslie, Danwei Huangfu

## Abstract

Human embryonic stem cells (hESCs) are notable for their ability to self-renew and to differentiate into all tissue types in the body. *NANOG* is a core regulator of hESC identity, and dynamic control of its expression is crucial to maintain the balance between self-renewal and differentiation. Transcriptional regulation depends on enhancers, but *NANOG* enhancers in hESCs are not well characterized. Here we report two *NANOG* enhancers discovered from a CRISPR interference screen in hESCs. Deletion of a single copy of either enhancer significantly reduced *NANOG* expression, compromising self-renewal and increasing differentiation propensity. Interestingly, these two *NANOG* enhancers are involved in a tandem duplication event found in certain primates including humans but not in mice. However, the duplicated counterparts do not regulate *NANOG* expression. This work expands our knowledge of functional enhancers in hESCs, and highlights the sensitivity of the hESC state to the dosage of core regulators and their enhancers.

## Introduction

Embryonic stem cells (ESCs) are characterized by two unique properties: self-renewal, the ability to proliferate while maintaining their original identity, and pluripotency, the potential to differentiate into all tissue types in the body. ESC identity is regulated by a core circuit of transcription factors that include NANOG^1,2^. *NANOG* is expressed in ESCs and down-regulated during differentiation^3,4^. This dynamic control of *NANOG* expression is crucial for the balance between the ESC and differentiated identities. Reducing *NANOG* expression compromised self-renewal, as evidenced by the decreased number of undifferentiated colonies and dropout from cell competition assays observed in mouse and human ESCs (mESCs and hESCs) with *NANOG* knockout or knockdown^5,6^. NANOG depletion also causes ESCs to mis-express markers for differentiated lineages, including extraembryonic endoderm, mesendoderm, and neuroectoderm^4,6–9^. Conversely, overexpression of *NANOG* in hESCs under differentiating conditions inhibits the commitment to neuroectoderm and definitive endoderm lineages^6,7,10^. Despite these insights from experimental manipulation of *NANOG* expression, the *cis*-regulatory mechanisms underlying its dynamic regulation in self-renewing and differentiating hESCs remain poorly understood.

Enhancers are a type of *cis*-regulatory element that ensures proper spatiotemporal regulation of gene expression. In mESCs, three enhancers have been shown to regulate the expression of *Nanog*, located 45kb upstream (−45kb), 5kb upstream (−5kb), and 9kb downstream (+9kb) of the *Nanog* transcriptional start site (TSS)^11–15^. Previous attempts to knockout the -5kb enhancer did not recover clones with biallelic deletions^13^, suggesting that this region is essential to mESC self-renewal. However, functional studies of *NANOG* enhancers in hESCs are lacking. Notably, mESCs and hESCs cultured under conventional media conditions correspond to distinct pluripotency states, with mESCs representing a more naïve state and hESCs a more primed state^16,17^. While both naïve and primed ESCs express core pluripotency regulators such as *NANOG*, *OCT4* (*POU5F1*), and *SOX2*, distinct enhancer usage has been reported for *OCT4* between these states, highlighting differences in the enhancer landscape^18–21^. This emphasizes the need to characterize *NANOG* enhancers specifically in primed hESCs.

In addition to cell state differences, the genome architecture of the *NANOG* locus also differs between human and mouse. A tandem duplication event involving a genomic region spanning from *NANOG* to approximately 200kb downstream is observed in certain primates, including humans. This results in two pairs of duplicated genes: *NANOG*/*NANOGP1*, and *SLC2A14/SLC2A3*^10,22–24^. The duplicated genes diverge in expression pattern and function. While *NANOG* is highly expressed and essential in both naïve and primed hESCs, *NANOGP1* is more selectively expressed in naïve hESCs and is dispensable for the maintenance of either naïve or primed pluripotency^23^. Both SLC2A3 and SLC2A14 belong to the glucose transporter (GLUT) family. *SLC2A3* is ubiquitously expressed, whereas *SLC2A14* expression is restricted to the testis^24^. Whether enhancers in this region are also duplicated and functionally divergent remains to be explored.

Enhancers have been defined based on different assays, ranging from biochemical annotation, episomal reporter activity, and gene expression changes upon perturbations in the endogenous genomic loci^25–27^. In order to discover functional enhancers, we adopted the last and most stringent approach, conducting an unbiased CRISPR interference (CRISPRi) screen in hESCs. Using dCas9-KRAB, we targeted putative enhancer regions of selected transcription factors, including *NANOG*, in their native genomic context. By using hESC self-renewal as a single, cell state-level readout, we were able to identify enhancers that regulate pluripotency through different target genes. The hits included two *NANOG* enhancers, named e1 and e2, consistent with the known requirement of *NANOG* for hESC self-renewal^6^. Both enhancers are duplicated in a region downstream of *NANOG* (flanking *NANOGP1*), but homozygous deletions of either duplicated enhancer (named e1′ and e2′) did not affect *NANOG* expression. In contrast, we were only able to generate heterozygous deletion lines for e1 and e2, suggesting they are essential for hESCs self-renewal. We show that *NANOG* expression is exquisitely sensitive to enhancer activity, as deleting even a single copy of either e1 or e2 significantly reduced its expression. Furthermore, the decreased *NANOG* expression resulting from the heterozygous enhancer deletions compromised hESC self-renewal and increased the propensity for differentiation. Our findings demonstrate that the *NANOG* enhancers regulate *NANOG* expression to maintain hESC identity.

## Results

### CRISPRi enhancer screen uncovers *NANOG* enhancers

In order to systematically identify functional enhancers in human pluripotency, we performed a CRISPRi screen in primed hESCs with inducible dCas9-KRAB. We applied a gRNA library^28^ tiling chromatin-accessible regions within 2Mb of the pluripotency regulators *NANOG*, *OCT4*, and *SOX2* (Figure 1A; Methods). We reasoned that upon induction of dCas9-KRAB expression, gRNAs targeting functional enhancers of genes that contribute to hESC self-renewal would be depleted over time. By applying an average gRNA |z-score| > 2 as a cutoff, we identified four hit regions associated with *OCT4* and *NANOG* (Figures 1B and 1C, Table S1-2). The hit regions associated with *OCT4* were the proximal enhancer (PE) and distal enhancer (DE), regulatory elements previously characterized in hESCs and conserved in mESCs^18–21^, thus validating our screening strategy (Figures S1A and S1B, Table S2).

**Figure 1.**
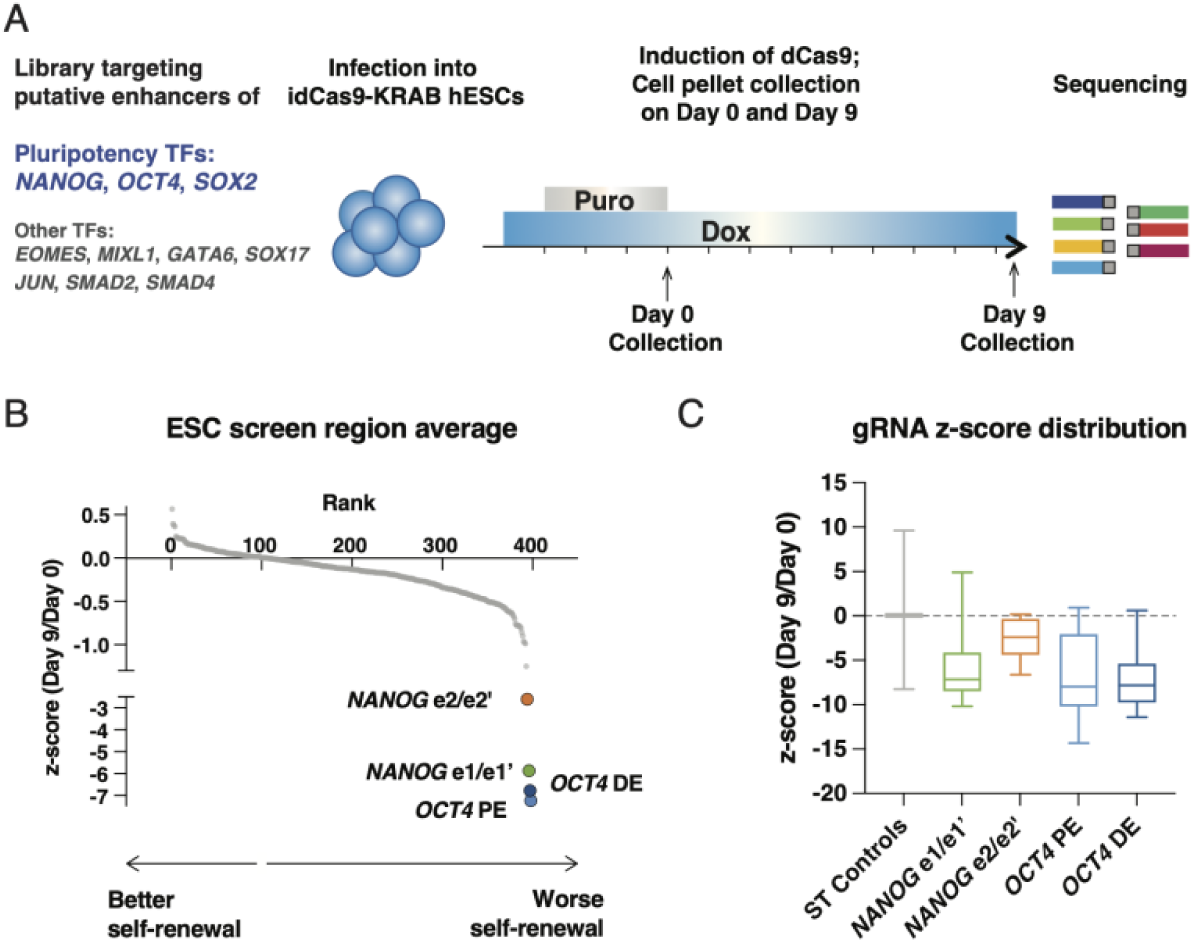
CRISPRi screen identified novel *NANOG* enhancers. A. Schematic of the enhancer screen in hESC. B. Results from the enhancer screen showing the average z scores of log_2_FC(Day 9 normalized reads/Day 0 normalized reads) of gRNAs targeting each region. Each dot represents a putative enhancer or the aggregate of safe-targeting (ST) gRNAs. Regions with average |z-score| > 2 are highlighted. C. Distribution of the z scores of ST gRNAs and the hit regions. The solid lines indicate the median values, the boxes contain 25^th^ to 75^th^ percentiles of data, and the whiskers mark maximum and minimum values.

In contrast to the known roles of the *OCT4* enhancers, the two hit regions associated with *NANOG* have not been functionally characterized before (Table S2). We named them *NANOG*-e1 and *NANOG*-e2. A closer inspection revealed that both *NANOG-*e1 and e2 are duplicated. *NANOG-*e1, 1.6kb upstream of *NANOG* TSS, shares 94% sequence identity with a downstream region designated e1′, 101.4kb downstream of *NANOG* TSS and 1.6kb upstream of *NANOGP1* TSS (Figure S1C, Table S2). *NANOG-*e2, 9.5kb downstream of *NANOG* TSS, shares 96% sequence identity with a downstream region hereby named e2′, 116kb downstream of *NANOG* TSS and 13.5kb downstream of *NANOGP1* TSS (Figure S1D, Table S2). Due to the high degree of sequence similarities, several gRNAs included in the screen targeted both the upstream and downstream regions. Because of this ambiguity, we merged the two duplicated regions as e1/e1′ and e2/e2′, respectively, when calling hits (Figures 1B and 1C).

### *NANOG* enhancers e1 and e2 regulate *NANOG* expression in *cis*

We examined the chromatin features at e1, e2, and their duplicated counterparts. *NANOG-*e1, e2, and e2′ have increased chromatin accessibility, H3K27ac, and H3K4me1 levels, features associated with active enhancers (Figures 2A and S1E). They were also bound by OCT4 and NANOG, suggesting they received regulatory inputs from these core TFs. While e1′ exhibited lower accessibility, it was also marked by H3K27ac, H3K4me1, and OCT4 binding. We called topologically associated domains (TADs) on published Hi-C data^28^ and found that the entire duplicated region is located within the same TAD containing the *NANOG* gene body (Figures S1E and S1F). This chromatin architecture suggests that all four regions could interact with the *NANOG* promoters and functionally regulate the expression of *NANOG*.

**Figure 2.**
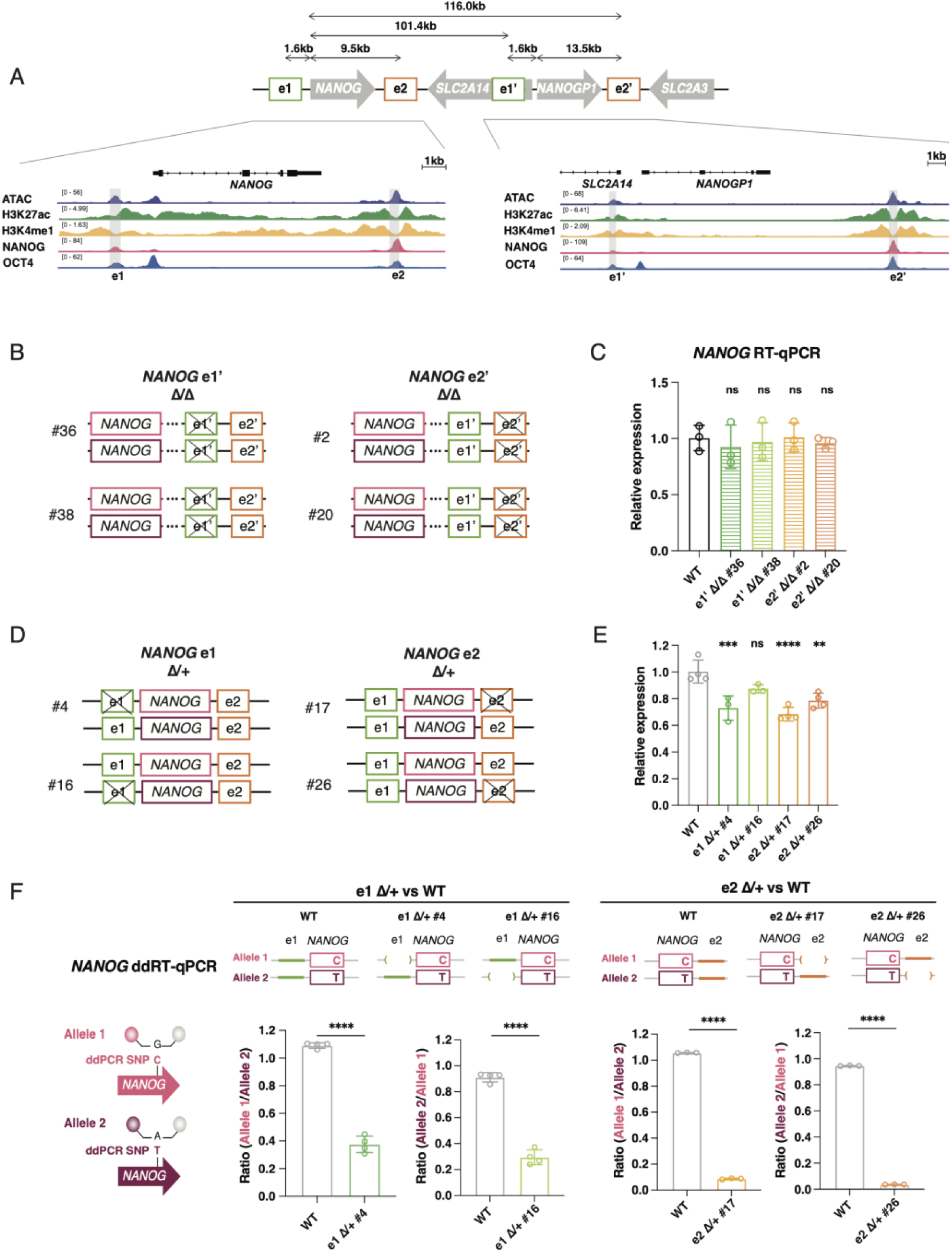
Validation of *NANOG*-e1 and e2 as functional enhancers. A. Schematics showing the duplicated region. Bottom left: ATAC and ChIP-seq tracks at the locus encompassing e1, *NANOG*, and e2; Bottom right: ATAC and ChIP-seq tracks at the *NANOGP1* locus encompassing the last exon of *SLC2A14*, e1 ′, *NANOGP1*, and e2 ′. ATAC, NANOG, and OCT4 ChIP-seq data have been published in Li et al.^44^, and histone ChIP-seq data have been published in Dixon et al.^45^. LiftOver was used to convert genome coordinates from hg19 to hg38. B. Schematic showing the genotypes of hESC lines with homozygous deletions (Δ/Δ) of e1′ or e2′. C. *NANOG* expression in e1′ or e2′ Δ/Δ lines measured by RT-qPCR. n = 3 biological replicates. D. Schematic showing the genotypes of hESC lines with heterozygous deletions (Δ/+) of e1 or e2. E. *NANOG* expression in e1 or e2 Δ/+ lines measured by RT-qPCR. n = 3 (for e1 Δ/+ lines) or 4 (for WT and e2 Δ/+ lines) biological replicates. F. Left: Schematic of the allele-specific probe design used in ddRT-qPCR; Right: Ratios of allele-specific *NANOG* expression as measured by ddRT-qPCR. The ratios in heterozygous enhancer lines are presented as that of the affected allele (*in cis* to the deleted enhancer) over the WT allele (*in cis* to the WT enhancer). The matching alleles were used to calculate the ratios in the WT line. n = 3 (for the e2 experiment) or 4 (for the e1 experiment) biological replicates, each performed in two ddRT-PCR technical replicates. Statistical analysis for C, E, and F: One-way analysis of variance (ANOVA) followed by Dunnett multiple comparisons test vs WT. ns p > 0.05; *p ≤ 0.05; **p ≤ 0.01; ***p ≤ 0.001; ****p ≤ 0.0001. Error bars indicate mean ± s.d.

To dissect the functional roles of e1, e2, and their duplicated counterparts, we performed targeted deletions in hESCs using paired gRNAs flanking each region. In all lines, we confirmed that the duplicated counterpart of the targeted regions remained intact (Table S3). We generated homozygous deletion lines for e1′ and e2′ (Figures 2B, S2A and S2B; Table S3), and RT-qPCR analysis revealed no change in *NANOG* expression (Figure 2C). In contrast, we were only able to recover heterozygous clones for e1 and e2 (e1 Δ/+ and e2 Δ/+), suggestive of their essential roles in hESC self-renewal (Figures 2D, S2C and S2D; Table S3). RT-qPCR identified a significant 20-30% reduction in *NANOG* expression in three of the four heterozygous enhancer deletion lines examined. A fourth line showed a similar trend, though this difference did not reach statistical significance (Figure 2E).

To address RT-qPCR variability and to quantitatively assess the *cis*-regulatory impact of enhancer deletions on *NANOG*, we conducted allele-specific digital droplet (dd) RT-qPCR using probes targeting a heterozygous SNP (named “ddPCR SNP”) in the *NANOG* transcript (Figure 2F, Table S4). We phased the WT allele and the enhancer deletion allele in relation to the ddPCR SNP alleles (Figures S2E and S2F, Methods). The two heterozygous lines for each enhancer had deletions *in cis* to different ddPCR SNP alleles, allowing us to control for potential expression variations between the two alleles. In WT hESCs, the two *NANOG* alleles were transcribed at a similar level. However, in all four enhancer Δ/+ lines, the *NANOG* allele *in cis* to the deleted enhancer was consistently transcribed at a significantly lower level than the allele *in cis* to the unedited enhancer (Figure 2F). Taken together, our results support that e1 and e2 regulate *NANOG* expression in an allele-specific manner, a hallmark of *bona fide* enhancers. In addition, the analysis of *cis* effect of enhancer deletions shows a loss of >55% (from the e1 deletion) and >90% (from the e2 deletion), resulting in an additive loss exceeding 100%. This “sub-subtractive” behavior in response to enhancer perturbations indicates that the endogenous e1 and e2 enhancers regulate *NANOG* in a super-additive manner, as also observed in other studies^29,30^, suggesting cooperative interactions between these enhancers.

### Heterozygous deletions of *NANOG* enhancers impair self-renewal and promote differentiation

To further investigate the cellular consequences of reduced *NANOG* expression through enhancer perturbations, we compared self-renewal and differentiation between WT and heterozygous enhancer lines. To assess hESC self-renewal, we labelled different lines with unique DNA barcodes^31^ and pooled them in a mixed culture. Barcode representation, used as a surrogate for relative cell growth and survival, was tracked by sequencing a fraction of the pooled cells every three days over a three-week period (Figure 3A; Methods). Both e1 Δ/+ and e2 Δ/+ lines exhibited a significant self-renewal disadvantage compared to WT lines (Figure 3B), supporting the requirement of these enhancers for hESC optimal self-renewal. This observation is also consistent with our CRISPRi screening results and our inability to recover homozygous deletion lines for e1 and e2. We then investigated the impact of enhancer perturbations on differentiation propensity by conducting definitive endoderm (DE) and neuroectoderm (NE) differentiation. We found that the e1 Δ/+ and e2 Δ/+ lines differentiated into DE and NE lineage more efficiently than WT, as indicated by the higher percentages of cells expressing the respective lineage marker (SOX17 for DE, and PAX6 for NE) (Figures 3C and 3D). This suggests that although *NANOG* expression is regulated by at least two enhancers, deletion of just one copy of either enhancer is sufficient to compromise hESC self-renewal and prime hESCs for differentiation.

**Figure 3.**
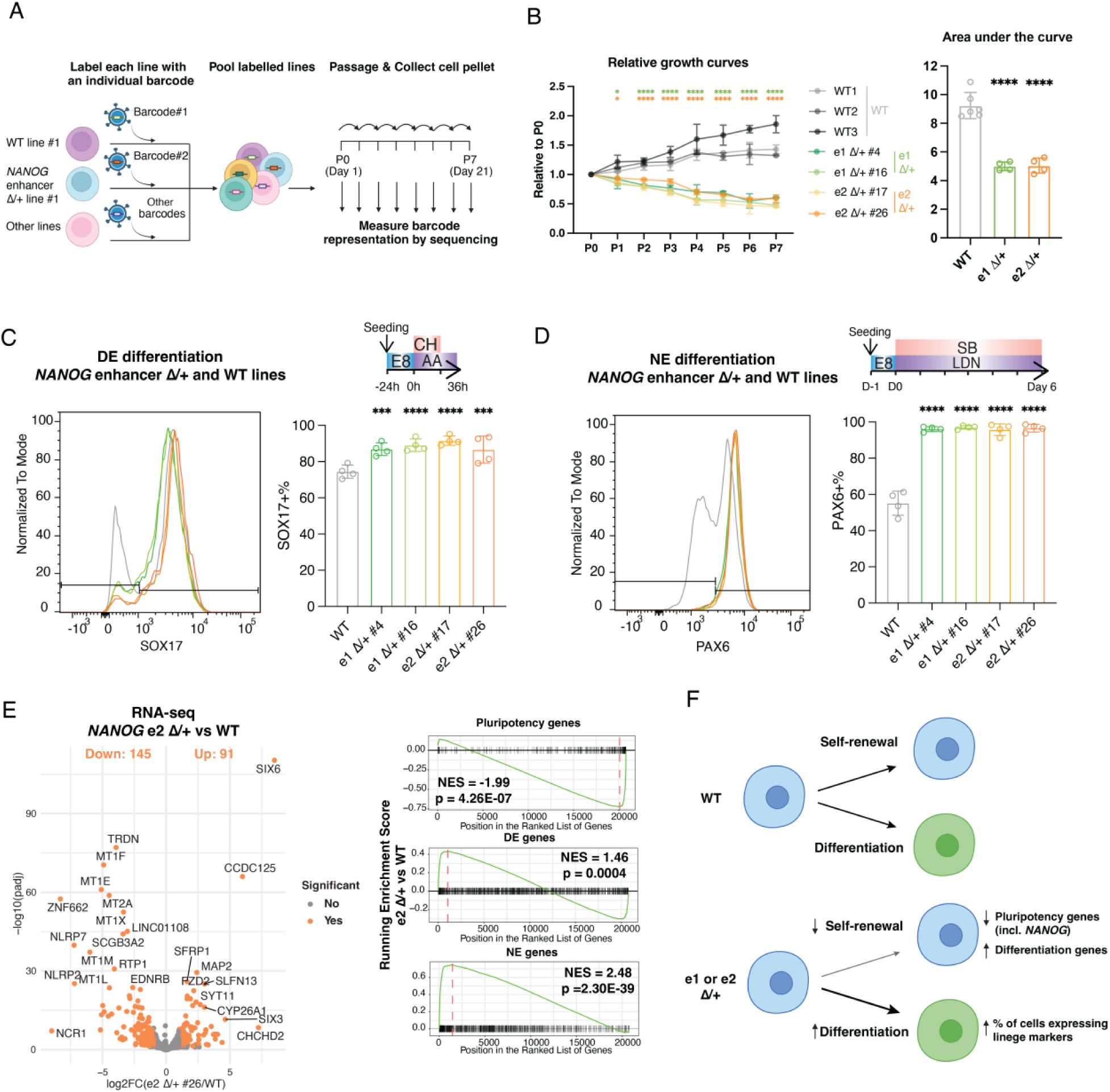
*NANOG* enhancers safeguard self-renewal and prevent differentiation. A. Schematic of the pooled self-renewal assay. *NANOG* Δ/+ enhancer lines gradually dropped out in the pooled self-renewal assay. Representation for each line was normalized to that at P0. For statistical analysis, lines with the same genotype were combined. n = 6 for WT (two replicates each for three lines) or 4 for e1 Δ/+ and e2 Δ/+ (two replicates each for two lines). Statistical analysis for the left panel: Two-way analysis of variance (ANOVA) followed by Šidák’s multiple comparisons test vs WT at each timepoint. Statistical analysis for the right panel: One-way analysis of variance (ANOVA) followed by Dunnett multiple comparisons test vs WT of the total area under the growth curve. ns p > 0.05; *p ≤ 0.05; **p ≤ 0.01; ***p ≤ 0.001; ****p ≤ 0.0001. Error bars indicate mean ± s.d. B. Flow cytometry showing increased DE differentiation efficiency of *NANOG-*e1 and e2 Δ/+ lines compared to WT cells. Left: histogram of SOX17 expression from one representative experiment; Right: summary data from 4 independent differentiations. C. Flow cytometry showing increased NE differentiation efficiency of *NANOG-*e1 Δ/+ and e2 Δ/+ lines compared to WT cells. Left: histogram of PAX6 expression from one representative experiment; Right: summary data from 4 independent differentiations. Statistical analysis for C-D: One-way analysis of variance (ANOVA) followed by Dunnett multiple comparisons test vs WT. ns p > 0.05; *p ≤ 0.05; **p ≤ 0.01; ***p ≤ 0.001; ****p ≤ 0.0001. Error bars indicate mean ± s.d. D. RNA-seq of a *NANOG-*e2 Δ/+ line and a WT line. n = 3 biological replicates, and GSEA for enrichment of pluripotency and differentiation gene sets ranked by expression log2FC in e2 Δ/+ vs WT. 91 up-regulated genes and 145 down-regulated genes were identified using |log2FC| > 1 and p.adj ⩽ 0.05 as cutoffs for significant differential gene expression and labelled in orange on the volcano plot. E. Schematic summarizing the phenotypes of *NANOG* enhancer Δ/+ lines in pluripotency and differentiation.

To investigate gene expression changes associated with these cellular phenotypes, we performed RNA-seq on a heterozygous enhancer line (e2 Δ/+) and a WT line at the hESC stage (Figure 3E). GO term analysis revealed up-regulation of genes associated with development, particularly those involved in the neural lineage, and down-regulation of metallothionein (MT) genes associated with metal homeostasis in the heterozygous enhancer line (Figures 3F and S3A). A previous study reported that MT genes were down-regulated during hESCs’ differentiation to definite endoderm^32^, consistent with our observation of increased differentiation propensity in enhancer heterozygous lines. To further explore the expression changes of genes involved in pluripotency and differentiation, we utilized our published RNA-seq of hESCs undergoing DE differentiation^28^ and performed an additional RNA-seq on NE differentiation. We defined genes up-regulated in DE and NE differentiation as DE or NE gene sets respectively, and genes down-regulated in both lineages as pluripotency genes (Figure S3B). GSEA analysis showed that DE and NE genes were significantly enriched for up-regulation in the e2 Δ/+ line, and pluripotency genes were enriched for down-regulation (Figure 3E). These results are consistent with previously observed effects of reduced *NANOG* expression in hESCs^6,7^. Together with the cellular phenotypes of the enhancer Δ/+ lines, they demonstrate that the *NANOG-*e1 and e2 enhancers are crucial for supporting full *NANOG* expression and maintaining the pluripotent identity of hESCs.

## Discussion

Here we report the identification of two *NANOG* enhancers, *NANOG*-e1 and *NANOG*-e2, through a CRISPRi enhancer screen. A region overlapping e1 was previously shown to bind PBX1 and drive luciferase reporter activity in hESCs^33^; however, no functional interrogation via deletion or perturbation of e1 or e2 at their endogenous loci has been performed. Our functional characterization of these two enhancers demonstrated that losing just one copy of either e1 or e2 substantially reduced *NANOG* expression and affected hESC self-renewal and pluripotency (Figure 3F). These results highlight the pluripotency state’s sensitivity to the dosage of its core regulators and the level of enhancer activity.

Different strategies have been employed to identify enhancers in hESCs. Using an episomal reporter assay^34^, tens of thousands of active enhancers were identified, creating rich datasets to study sequence features contributing to enhancer activity, but linking these candidate enhancers to their target genes remains challenging. Examination of this screen data revealed that genomic regions overlapping *NANOG*-e1 and e2 were classified as active enhancers and therefore corroborated our findings. Reporter assays are highly scalable but do not assess the roles for enhancers in their native genomic contexts. Advances in CRISPR technology make it possible to functionally characterize enhancers through perturbing them in their endogenous chromatin context. Multiple *OCT4* enhancers have been discovered through Cas9-mediated screens, using *OCT4*-GFP reporter expression as a direct readout^35,36^. Moving beyond the interrogation of enhancers for a single gene, combining CRISPRi perturbation with scRNA-seq has enabled assessment of enhancer perturbation effect on many target genes at the same time in cancer cell lines^37,38^. While powerful, it remains challenging and resource-intensive to conduct such screens. Nevertheless, they are beginning to show promise in hESC systems. For instance, a recent study successfully interrogated 25 putative enhancers in hESC cardiac differentiation^39^. In our study, we combined CRISPRi perturbation with a cell state-level readout, which enabled us to interrogate functional enhancers of multiple genes based on their impact on hESC self-renewal. We have also applied this approach in differentiation contexts in other works. By using the expression of key lineage markers as a readout, we have successfully identified enhancers not only for these marker genes but also for other genes that regulate the acquisition of the differentiated identity^28,40^. While this strategy is cost-effective and relatively robust, it is not optimal for identifying enhancers of multiple target genes that do not converge on a single readout. Given the current challenges, we feel that different strategies should be considered for enhancer discovery, balancing sensitivity, scalability, and cost.

The requirement for both *NANOG* enhancers in primed hESCs contrasts with the findings in mouse. Conversion of genome coordinates by LiftOver^41,42^ shows that *NANOG-*e1 corresponds to the -5kb *Nanog* enhancer in mouse, and *NANOG-*e2 corresponds to the +9kb *Nanog* enhancer in mouse, but BLAST alignment^43^ identified little sequence identity between the human and mouse enhancers (Figure S4, Table S5). In comparison, the *OCT4*-DE and PE enhancers show much greater sequence conservation between mice and humans (Figure S4, Table S5). Previous study attempting to knockout the -5kb *Nanog* enhancer in mESCs did not recover clones with biallelic deletions, but found that a monoallelic deletion decreased *Nanog* expression by ∼50%, suggesting this enhancer was essential for *Nanog* expression and mESC self-renewal^13^. No deletion experiment of the +9kb enhancer has been done in mESCs, but it is shown to drive transcription in plasmid reporter assays^11^. In a CRISPR-Cas9-mediated screen of regulatory elements in mESCs, gRNAs targeting the +9kb region were associated with decreased *Nanog* reporter signals, although not to the same extent as those targeting the -5kb enhancer^15^. Taken together, these studies suggest that the -5kb enhancer is the predominant one in mESCs, and the +9kb enhancer may help finetune *Nanog* expression. In comparison, both *NANOG*-e1 and e2 are essential for hESC self-renewal. As mESCs represent a more naïve state of pluripotency, it will be of interest to explore whether hESCs preferentially rely on one of the *NANOG* enhancers in the naïve state. In addition, *NANOGP1* is selectively expressed in naïve but not primed hESCs^23^, raising the question of whether the duplicated enhancers e1′ and e2′ regulate this duplicated gene and exhibit stage-specific regulatory activity. Investigating these questions will not only expand our knowledge of the regulatory mechanisms governing different states of human pluripotency, but also shed light on the evolutionary conservation and diversification of enhancer functions.

---

**Figure S1.**
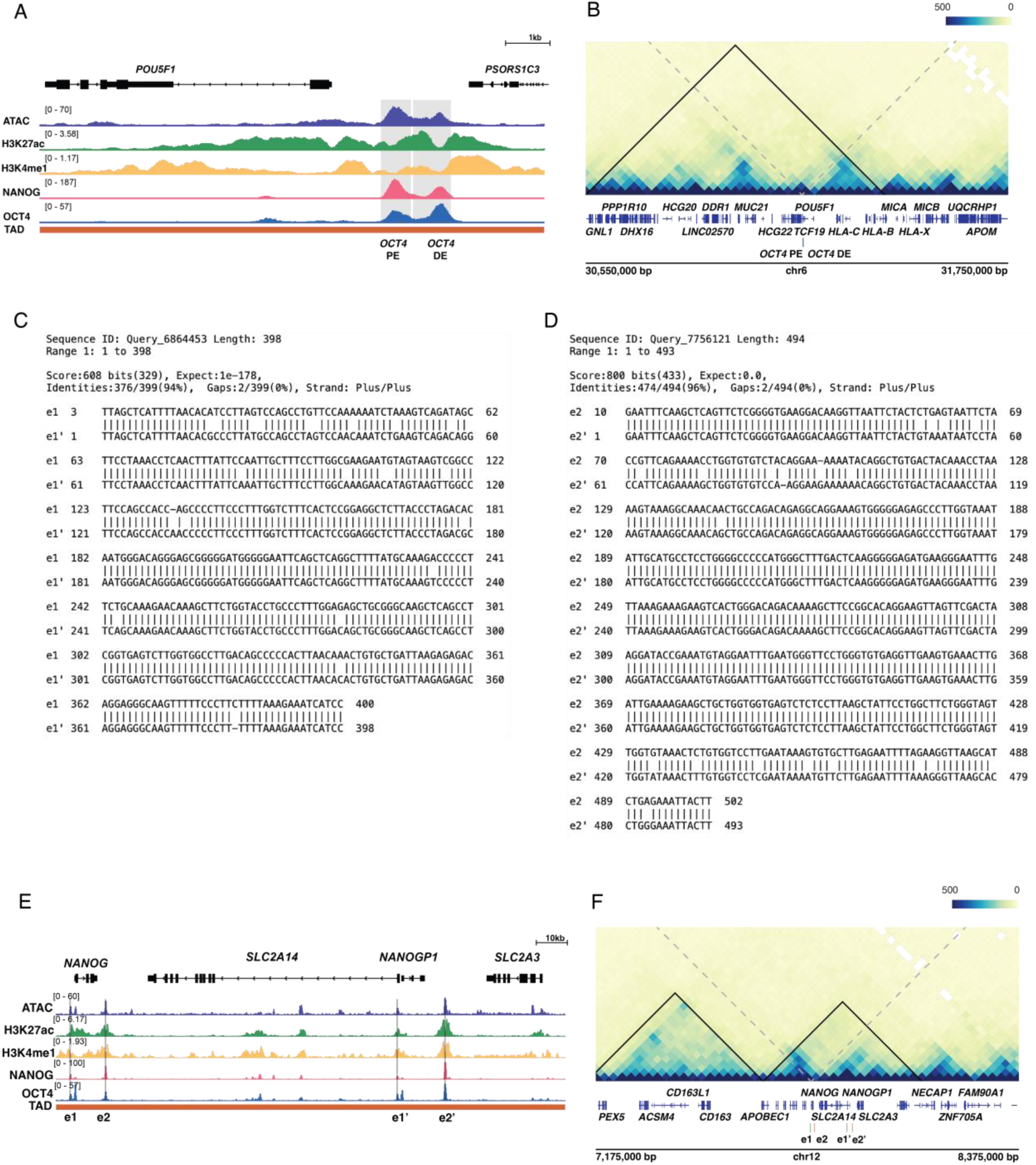
Chromatin modifications and architecture at enhancer hit regions. A. ATAC, ChIP-seq, and Hi-C TAD at the *OCT4* (*POU5F1*) PE/DE locus. B. Hi-C normalized contact frequency showing the 1.2Mb genomic window encompassing *OCT4* (*POU5F1*). Solid black lines indicate TAD boundaries, and grey dashed lines indicate interactions with *OCT4* promoter. C. BLAST alignment of *NANOG-*e1 and e1′ showing significant sequence similarity. D. BLAST alignment of *NANOG-*e2 and e2′ showing significant sequence similarity. E. ATAC, ChIP-seq, and Hi-C TAD at the entire duplicated region encompassing *NANOG*. F. Hi-C normalized contact frequency showing the 1.2Mb genomic window encompassing *NANOG.* Solid black lines indicate TAD boundaries, and grey dashed lines indicate interactions with *NANOG* promoter. ATAC, NANOG, and OCT4 ChIP-seq data have been published in Li et al.^44^, and histone ChIP-seq data have been published in Dixon et al.^45^. Hi-C data have been published in Luo et al.^28^. For sequencing data aligned to hg19, LiftOver was used to convert genome coordinates to hg38.

**Figure S2.**
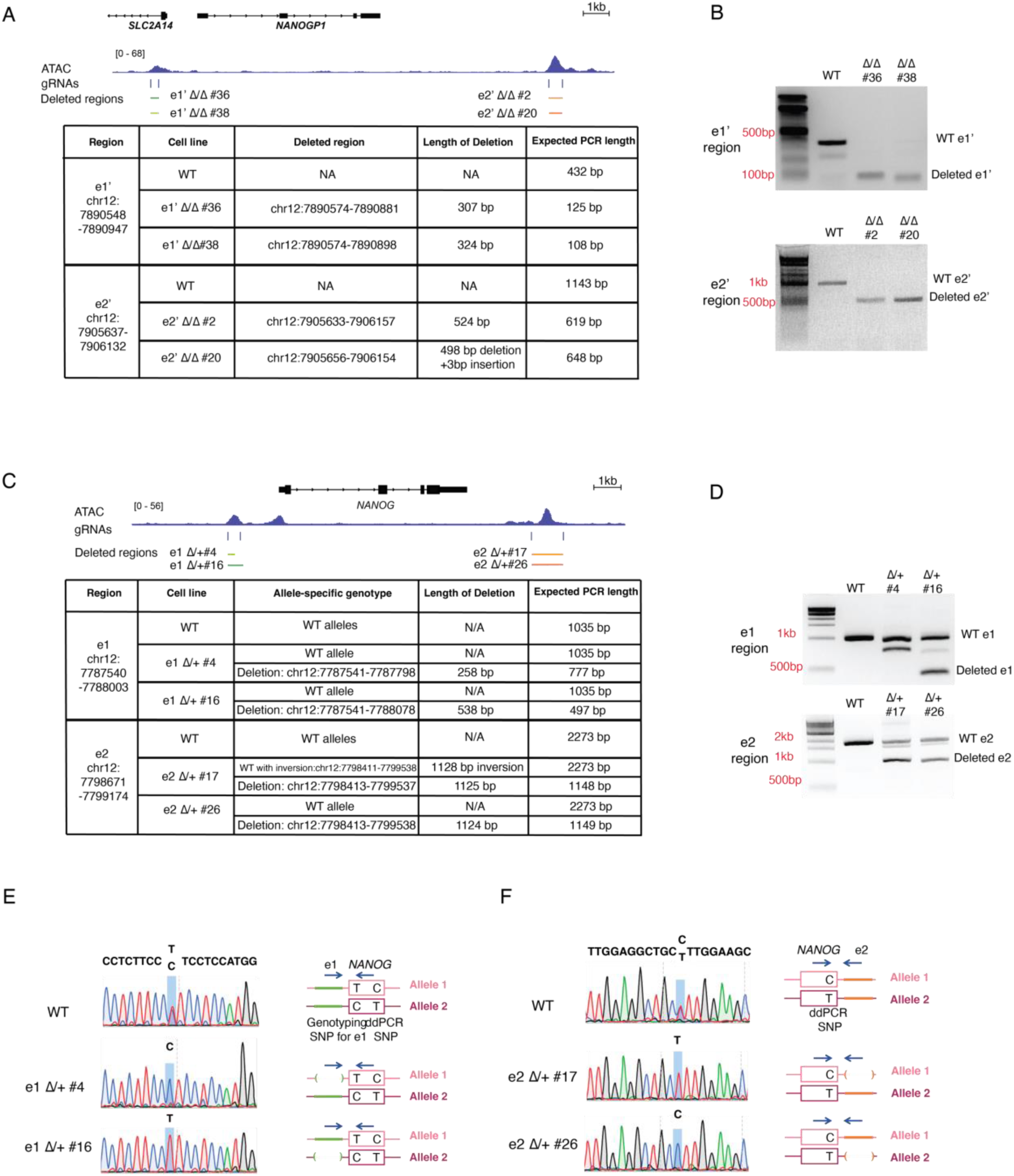
Genotypes of enhancer deletions and ddPCR SNP. A. Gene tracks and table showing the locations of e1 ′ and e2 ′ deletions in Δ/Δ lines. B. Top: gel picture showing PCR genotyping of the e1′ region in WT and e1′ Δ/Δ lines; bottom: gel picture showing PCR genotyping of the e2′ region in WT and e2′ Δ/Δ lines. C. Gene tracks and table showing the locations of e1 and e2 deletions in Δ/+ lines. In e2 Δ/+ #17, the sequence of the remaining e2 was inverted, but *NANOG* transcription from this allele was comparable to WT. D. Top: gel picture showing PCR genotyping of the e1 region in WT and e1 Δ/+ lines; bottom: gel picture showing PCR genotyping of the e2 region in WT and e2 Δ/+ lines. E. Sequencing spectrogram of the amplicon partially covering intact e1 and the genotyping SNP for e1 to phase WT and deleted e1 in relation to the ddPCR SNP. F. Sequencing spectrogram of the amplicon partially covering intact e2 and the ddPCR SNP to phase WT and deleted e2 in relation to the ddPCR SNP.

**Figure S3.**
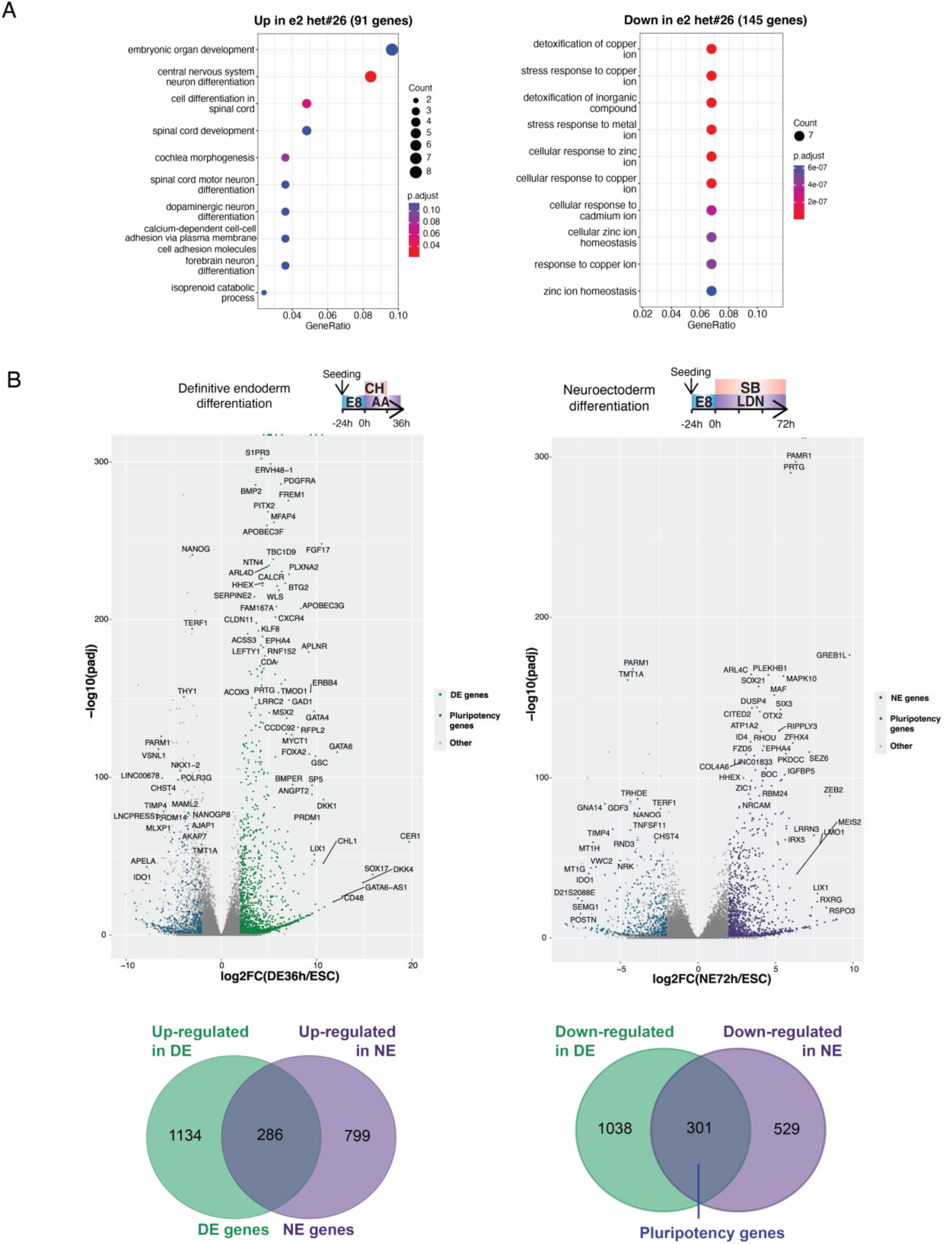
RNA-seq in hESCs with heterozygous enhancer deletion and in differentiation. A. Gene Ontology analysis of biological processes enriched in genes up-regulated (left; n = 91) or down-regulated (right; n = 145) in e2 Δ/+ compared to WT hESCs. Cutoffs for significant differential gene expression: |log2FC| > 1 and p.adj ≤ 0.05. B. Definition of DE and NE genes based on differential gene expression at DE 36h and NE 72h compared to hESCs respectively. Cutoffs for significant differential gene expression: |log2FC| > 2 and p.adj ≤ 0.05. Gene set definition is indicated in the Venn diagrams.

**Figure S4.**
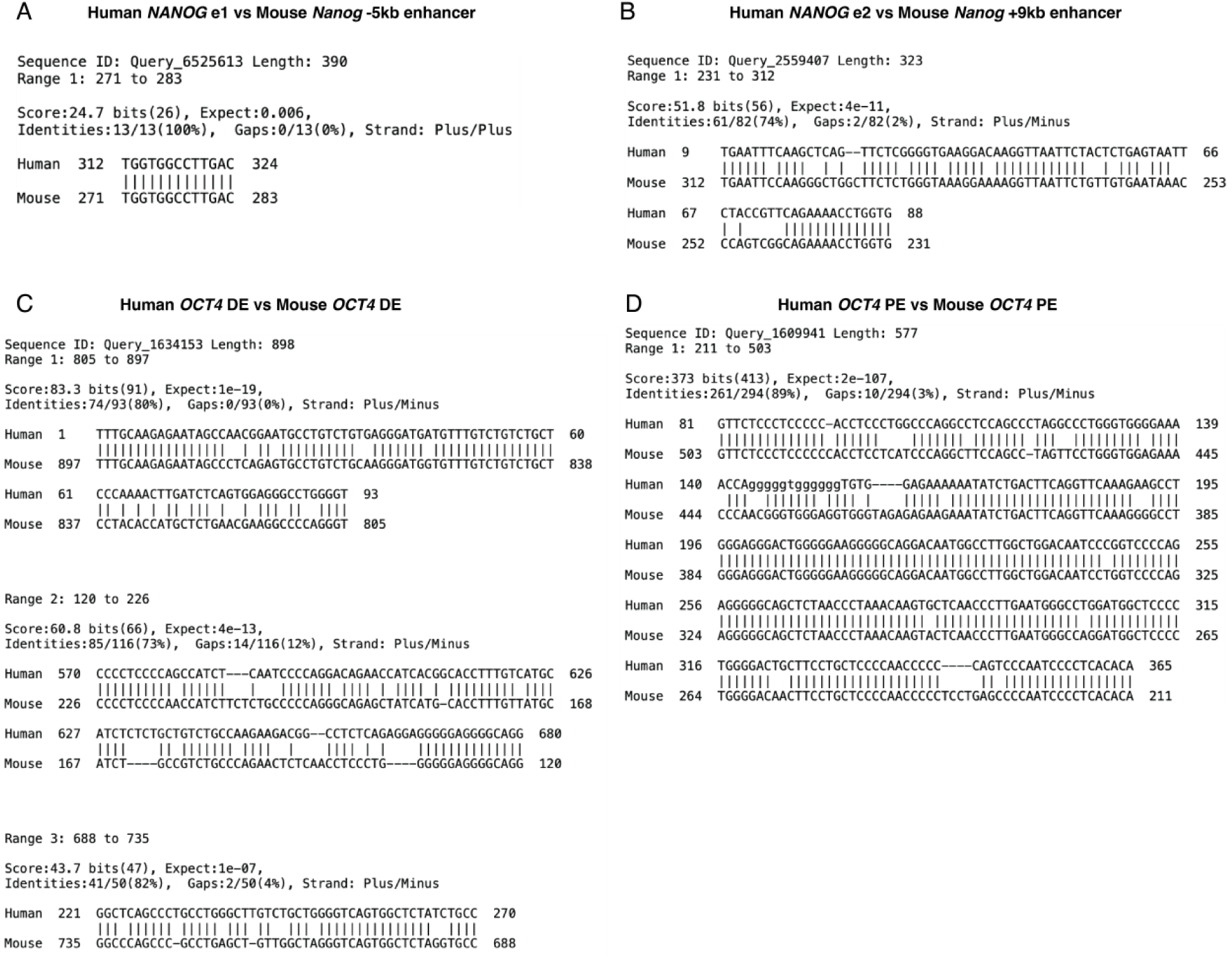
Comparing sequence identity of pluripotency enhancers in human and mouse. A. BLAST alignment of human *NANOG-*e1 and mouse *Nanog* -5kb enhancer. B. BLAST alignment of human *NANOG-*e2 and mouse *Nanog* +9kb enhancer. C. BLAST alignment of human *OCT4-*DE and mouse *Oct4-*DE. D. BLAST alignment of human *OCT4-*PE and mouse *Oct4-*PE.

## Author contributions

J.Y. and D.H. conceptualized the study, devised the experiments, and interpreted results. J.Y. performed most of the experiments and analyzed the results. R.L. performed the CRISPRi screen. B.P.R. performed the NE differentiation and RNA-seq. D.L. assisted with the cell growth competition assay. W.W. visualized Hi-C data and called TADs and was supervised by C.S.L.. J.Y. and D.H. wrote the manuscript; all authors provided editorial advice.

## AI-assisted technologies in the writing process

During the preparation of this work the authors used ChatGPT developed by OpenAI to improve the clarity of some sentences. After using this tool, the authors reviewed and edited the content and take full responsibility for the content of the publication.

## Acknowledgements

We acknowledge the assistance from the following MSKCC Cores: Antibody & Bioresource, Flow Cytometry, Gene Editing & Screening, Integrated Genomics Operation (IGO) and Molecular Cytogenetics. We thank R. Garippa, H. Liu and S. Mehta for assistance with CRISPR library generation and HiSeq for quantifying gRNA abundance after CRISPRi screen, and Dr. Andrea Farina for designing and executing the ddPCR assays. We thank R. Hosuru and S. Adeniyi for assisting with experiments not included in the manuscript, S. Kaplan, T. Guo, and J. Pulecio for editorial input, and A.-K. Hadjantonakis, L. Studer, and T. Norman for insightful advice. This study was funded in part by the National Institutes of Health grant U01HG012051 (D.H.), National Institutes of Health grant U01DK128852 (C.S.L., D.H.), Starr Tri-I Stem Cell Initiative #2019-001 (D.H.), NIH/NCI MSKCC Cancer Center Support Grant P30CA008748, a Beatrice P. K. Palestin Fellowship and a Bruce Charles Forbes Fellowship (to D.L.), an NIH T32 Training Grant (T32GM008539 to B.P.R.), Cycle for Survival (to IGO), and the Marie-Josée and Henry R. Kravis Center for Molecular Oncology (to IGO).

## Data availability

New sequencing data from this study (RNA-seq of e2 heterozygous line, WT hESC, and WT differentiation at NE72h) have been deposited in the GEO database under accession code GSE283612. The ATAC-seq, OCT4 ChIP-seq, and NANOG ChIP-seq data have been previously published under accession code GSE109524. The H3K27ac and H3K4me1 ChIP-seq data have been previously published under accession code GSE150072. The RNA-seq data at DE 0h and 72h and the Hi-C data have been previously published under accession code GSE213394.

## Methods

### Culture of hESCs

H1 (NIHhESC-10-0043) and HUES8 (NIHhESC-09-0021) hESC lines are from male donors with no reported disease information. All experiments were conducted per NIH guidelines and approved by the Tri-SCI Embryonic Stem Cell Research Oversight (ESCRO) Committee. Cell lines were authenticated by the standard short tandem repeat (STR) profiling using the Memorial Sloan Kettering Cancer Center (MSKCC) Integrated Genomics Operation core facility, and cells were regularly confirmed to be mycoplasma-free by the Antibody & Bioresource Core Facility. No bacterial or fungal contamination was detected during data collection.

hESCs were maintained in Essential 8 medium (E8; Thermo Fisher Scientific, A1517001) on tissue culture-treated polystyrene plates coated with 5 μg/mL vitronectin (Thermo Fisher Scientific, A14700) at 37 °C with 5% CO2. For regular maintenance, cells were passaged every 2-4 days, where they were treated with 0.5 mM EDTA (KD Medical, RGE-3130) in PBS for dissociation. For long term preservation, cells were dissociated, resuspended in E8 medium with 10% DMSO (Santa Cruz Biotechnology, sc-358801), and frozen in liquid nitrogen. For seeding cells, 10 μM Rho-associated protein kinase (ROCK) inhibitor Y-27632 (Selleck Chemicals, S1049) was added into the culture medium. Cell counting was performed on a Vi-CELL XR Cell Viability Analyzer (Beckman Coulter). We used idCas9–KRAB SOX17^eGFP/+^ HUES8 line^28^ for experiments involving inducible dCas9-KRAB expression, and the H1 OCT4^GFP/+^ iCas9 line^46^ for enhancer deletion experiments. New cell lines were derived from cryopreserved stocks within 4–10 passages and used for experiments within 10 subsequent passages.

### CRISPR enhancer screen

The gRNA library was generated for a previous study^28^. The library targets 394 putative enhancer regions and includes 1100 safe-targeting gRNAs^47^. The enhancer screen in ESCs was performed concurrently with the DE screen in Luo et al. with slight modifications. A total of 35 million idCas9–KRAB SOX17^eGFP/+^ HUES8 hESCs were infected with the lentiviral gRNA library at a low MOI of ∼0.3 with a targeted coverage of ∼1,000 fold on day 0 in 15-cm plates. A total of 6 μg/ml protamine sulfate per plate was added during the first 24 h of infection. One day after infection, cells were treated with 2 μg/ml doxycycline to induce dCas9–KRAB expression, which continued till the end of the screen. Infected cells were selected with 1 μg/ml puromycin from day 1 to day 3. After selection, 20M cells were collected to extract Day 0 genomic DNA and 12M cells were seeded into 4 15-cm plates (3M/plate) for maintenance culture. 12M cells were passaged onto new plates every 3 days for 3 passages for the final collection of 20M Day 9 cells. Genomic DNA from both Day 0 and Day 9was extracted using the QIAGEN Blood and Cell Culture DNA Maxi Kit and quantified by Qubit (Thermo Fisher Scientific, Q32850) following the manufacturer’s guidelines. An amount of genomic DNA representing the gRNA library for 1000x from each sample was amplified. Amplicons were quantified by Qubit and Bioanalyzer (Agilent) and sequenced on the Illumina HiSeq 2500 platform by the MSKCC Integrated Genomics Operation core.

### Data analysis for the CRISPRi enhancer screen

The analysis for the core enhancer screen was performed as previously described^28^ with slight modifications. After aligning the HiSeq reads to the gRNA library reference sequences, the counts for each gRNA were obtained and normalized to the total reads of each sample. The normalized counts for a gRNA in the Day 0 and Day 9 samples were averaged and logged by a base of 2 to derive the log_2_(Average of Abundance). Then, the log_2_ fold change (FC) of normalized counts for each gRNA in Day 9 over Day 0 was calculated. A z-score was calculated for each gRNA by subtracting the mean log_2_FC of all safe-targeting gRNAs from the gRNA’s log_2_FC and dividing the difference by the standard deviation of log_2_FC of all safe-targeting gRNAs. All gRNAs were included in the analysis to retain hits, and gRNAs targeting multiple regions were included in the calculation for each region.

### Definitive endoderm differentiation

Definitive endoderm differentiation was performed as previously described^28,44^. Briefly, hESCs grown to ∼80% confluency in a “recovery passage” were washed with PBS and treated with TrypLE Select at room temperature for 3 min. After the treatment, TrypLE was removed, and hESCs were dissociated into single cells, resuspended in E8 medium, spun at 200g for 2 minutes, and resuspended in E8 medium with 10 μM ROCK inhibitor. 1 million cells were seeded per well on a 6-well plate (Fisher Scientific 0720080), or 0.3 million cells per well on a 12-well plate (Fisher Scientific, 0720082), coated with 5 μg/mL vitronectin. 24 hours after seeding, cells were washed with PBS and exposed to S1/2 medium containing 50 ng/mL Activin A (Bon-Opus Biosciences, C687-1MG) and 5 μM CHIR99021 (Tocris Bioscience, 4423). 24 hours later, the medium was changed to S1/2 containing only 50 ng/mL Activin A. After another 12 hours, cells were harvested for DE 36h time point. To make S1/S2 medium, MCDB131 medium (Thermo Fisher Scientific, 10372019) was supplemented with 1.5 g/L sodium bicarbonate (Research Products International, S22060), 1x Glutamax (Thermo Fisher Scientific, 35050061), 10 mM glucose (Sigma-Aldrich, G8769), and 0.5% BSA (LAMPIRE, 7500804).

### Neuroectoderm differentiation

Neuroectoderm differentiation was performed as previously described^46,48,49^. Briefly, confluent hESC culture was washed with PBS and treated with TrypLE Select (Thermo Fisher Scientific, 12563-029) at room temperature for 3 min. After the treatment, TrypLE was removed, and hESCs were dissociated into single cells, resuspended in E8 medium, spun at 200g for 2 minutes, and resuspended in E8 medium with 10 μM ROCK inhibitor. 0.2 million cells were seeded per well on a 12-well plate coated with 5 μg/mL vitronectin. 24 hours after seeding, cells were washed with PBS and exposed to Essential 6 medium (E6; Thermo Fisher Scientific, A1516401) containing 10 µM SB431542 (Tocris, 161410) and 500 nm LDN193189 (Cedarlane Labs, 04-0074-02). Then cells were changed into medium with the same formula every 24 hours for a total of 6 days and harvested.

### Hi-C analysis

Hi-C contact frequency was quantified and visualized as previously described^40^. Briefly, HiC-Pro^50^ (3.1.0) was used to align the individual Hi-C replicates to GRCh38, GCA_000001405.15, with alternative contigs removed, and then final merged libraries were obtained by combining all *.allValidPairs files where duplicates across biological replicates are maintained. For the contact map, normalized contact frequency was visualized using KR balancing at 25 kB resolution. TADs were called as previously described, using TopDom 0.10.1 via the interface available in Hi-C-DC+ with ICE balancing. TAD calling was performed using 50 kb bins, scanning 10 kb up and downstream to compute boundaries^51–53^ (Table S6).

### Enhancer deletion

Generation of heterozygous enhancer lines was performed as previously described^28,40^ with modifications. The enhancer to be deleted was targeted by one crRNA at each end and tracrRNA, both ordered from IDT and diluted in nuclease-free duplex buffer (11-01-03-01). H1 OCT4^GFP/+^ iCas9 hESCs were treated with 2 μg/mL doxycycline (Sigma Aldrich, D9891) 24 hours before, on the day of, and 24 hours after gRNA transfection to induce Cas9 expression. Cells were dissociated with TrypLE Select and seeded at 150,000 cells/well density in a vitronectin-coated 24-well plate in E8 medium containing 10 μM ROCK inhibitor. 7.5 pmol of each crRNAs and 15 pmol of tracrRNA were diluted in 25 μL Opti-MEM (Thermo, 31985070), and 1.5 μL Lipofectaine RNAiMax Transfection Reagent (Thermo, 13778100) was diluted separately in another 25 μL Opti-MEM. The two solutions were mixed together, incubated for 15 minutes at room temperature, and added drop-wise to freshly seeded cells. When transfected cells reached confluency, they were dissociated into single cells using TrypLE Select and seeded at 2000 cells/plate density on a vitronectin-coated 100 mm tissue culture dish (Millipore Sigma, CLS430167) in E8 medium containing 10 μM ROCK inhibitor for colony formation. A week after seeding, colonies were picked into individual wells on a vitronectin-coated 96-well plate and cultured until confluency. gDNA was extracted from each confluent 96-well (∼50,000 cells) by overnight incubation at 55C of cell lysate from the first passage with 1X PCR buffer (Sigma Aldrich, P2192) and 1 mg/mL proteinase K (Fisher Scientific, BP1700) and used for initial PCR genotyping. Enhancer heterozygous lines were expanded, split for the second-round gDNA collection, and cyro-preserved. Before thawing for use in experiments, the lines’ genotypes were further confirmed with PCR done on gDNA extracted from a 6-well (1-2 million cells) during the second-round gDNA collection by the QIAGEN Blood & Cell Culture DNA Maxi Kit (QIAGEN, 13362) and targeted sequencing (Eton Bioscience).TOPO cloning (Thermo Fisher Scientific, 450245) was further performed using the genotyping PCR product to confirm the genotype on both alleles and the homogeneity of genotype within the same culture. crRNA sequences and genotyping primers are listed in Tables S7 and S8.

### RNA isolation, reverse transcription and RT-qPCR

Total RNA was isolated from cell pellets using Quick-RNA Miniprep kits (ZYMO Research, R1055) per the manufacturer’s instructions, including the digestion and removal of DNA. Reverse transcription was performed using the High-Capacity cDNA Reverse Transcription Kit (Applied Biosystems, 4368814) with the addition of 20 units RNase inhibitor (Thermo Scientific, AM2696) on a Bio-Rad T100 thermocycler. 20 ng cDNA (measured as RNA before reverse transcription on a Nanodrop 2000 spectrophotometer, Thermo Fisher Scientific), 1x SYBR Green Master Mix (Thermo Fisher Scientific, A25776), and 0.5 μM each of forward and reverse primers were combined to make a 10 μL reaction mixture, and loaded onto a well on a MicroAmp Optical 384 well Reaction Plate (Thermo Fisher Scientific, 4309849), sealed with a MicroAmp Optical Adhesive Film (Thermo Fisher Scientific, 4360954). The measurement was performed on a QuantStudio 6 Flex Real-Time PCR System (Applied Biosystems) and analyzed using the 2^−ΔΔCt^ method. RT-qPCR primers are listed in Table S8.

### Genotyping allele-specific SNPs

To identify allele-specific SNPs on the *NANOG* transcript, total RNA was isolated from H1 hESCs and reverse transcribed into full-length cDNA with SuperScript™ III First-Strand Synthesis System (Thermo Fisher Scientific, 18080051). The *NANOG* transcripts were PCR amplified, inserted by In-Fusion (Takara Bio, 639648) into a plasmid (addgene: 83481) digested with NheI (NEB R0131) and FseI (NEB R0588S), and transformed into competent *E.coli* (NEB, C2987H). Out of the five SNPs identified to be heterozygous on the *NANOG* transcript in H1 hESCs from sequencing 8 individual clones, rs4012937 (“ddPCR SNP”) was chosen for the design of the allele-specific RT-ddPCR assay (Table S4).

The genotyping strategy for allele-specific SNPs is schematized in Figures S2E and S2F. To determine the genotype at the ddPCR SNP *in cis* to the WT e2 in e2 heterozygous lines, the region spanning the ddPCR SNP and part of the intact e2 was amplified using LongAmp Taq Polymerase (NEB, M0323S) and sequenced (Eton Bioscience). The genotype at the ddPCR SNP *in cis* to the deleted e2 was inferred thereafter. Since the genomic distance between the ddPCR SNP and e1 was challenging for PCR amplification, the genotype of rs4294629 (“Genotyping SNP” for e1), the heterozygous SNP located closest to the 5’ of the *NANOG* transcript, *in cis* to WT e1 was determined instead for e1 heterozygous lines. The region spanning the Genotyping SNP and part of the intact e1 was amplified using LongAmp Taq Polymerase and sequenced. The genotype at the Genotyping SNP *in cis* to the deleted e1 was inferred thereafter. The genotype of the ddPCR SNP *in cis* to WT and deleted e1 was inferred from the knowledge that allele T at the Genotyping SNP is *in cis* to allele C at the ddPCR SNP, and allele C at the Genotyping SNP is *in cis* to allele T at the ddPCR SNP in H1 hESCs, as determined from the cloning experiment (Supplementary Table 4, Supplementary Data 2). The primers for cDNA amplification, In-Fusion cloning, and SNP genotyping are listed in Table S8.

### Digital Droplet PCR (ddPCR)

The ddPCR assay was designed and performed by the Integrated Genomics Operation core at MSKCC as previously described with slight modifications^40^. Probes specific for allele C at the ddPCR SNP labelled with the HEX fluorophore and specific for allele T labelled with the FAM fluorophore and primers amplifying the region spanning the ddPCR SNP on the *NANOG* transcript were ordered from BioRad. RNA isolation and reverse transcription were performed as described in previous sections. Each ddPCR reaction consisted of 1 ng cDNA/reaction in duplicate (total 2 ng), 900 nM primers, 250 nM probes, and 1X ddPCR Supermix for probes (no dUTP, BioRad). ddPCR droplets were generated using the QX200 Droplet Generator (BioRad, 1864002). Droplets were read using QX200 Droplet Reader (BioRad, 1864003) and analyzed using the QuantaSoft Software (BioRad, 1864011). The primer and probe sequences are listed in Table S8.

### Flow cytometry

Flow cytometry was performed as previously described^28^. Briefly, for live GFP reporter flow cytometry, cells were dissociated with TrypLE Select, washed in FACS buffer (PBS with 5% Fetal Bovine Serum, Sigma Aldrich 12103C and 5 mM EDTA), stained with antibodies diluted in FACS buffer for 15 minutes, and then with DAPI (Sigma Aldrich, 32670) diluted in FACS buffer at 1 μg/mL for 5 minutes. For fixed cell flow cytometry, cells were dissociated, washed in FACS buffer, and incubated with LIVE-DEAD Fixable Violet Dead Cell Stain (Invitrogen, L34955) diluted per manufacturer’s instructions in FACS buffer for 15 minutes, and fixed with Fixation/Permeabilization concentrate diluted 1:3 in corresponding diluent (Fisher Scientific, 00522356/00512343) for 1 hour. Afterwards, cells were incubated with primary and then secondary antibodies diluted in 1x Permeabilization Buffer (Thermo Fisher Scientific, 00833356) for 1 hour each, and resuspended in FACS buffer. Flow cytometry data were collected using BD LSRFortessa or BD LSRII with BD FACSDIVA. Flow cytometry analysis and figures were generated in FlowJo v10. Flow antibodies are listed in Table S9.

### Growth competition assay

Growth competition assay was performed as described previously^46^. Briefly, individual LARRY barcode constructs were cloned from the LARRY barcode library^31^ (Addgene:140024), packaged into lentivirus, and infected into different cell lines so that each line was labelled with a unique barcode. A week after infection, GFP^high^ cells were selected by fluorescence-activated cell sorting (FACS) and cultured to confluency as individual clones. Then each cell line was dissociated, counted, and mixed in equal number to form a pool (100,000 cells per line), 200,000 cells of which were seeded and the rest harvested for collection of P0 genomic DNA. Cells were passaged by TrypLE Select dissociation every 3 days for a total of 21 days. At each passage, 200,000 cells were seeded and the rest harvested for genomic DNA extraction by the QIAGEN Blood & Cell Culture DNA Maxi Kit. LARRY barcodes were amplified by PCR using Q5 polymerase (NEB, M0491S; primers listed in TABLE), cleaned up using AmPure beads (Beckman Coulter, A63881), and sequenced for 5-10 million PE150 reads per sample by the MSKCC Integrated Genomics Operation core. CRISPResso2 (http://crispresso.pinellolab.org/submission) was used for demultiplexing and quantification of barcode representation. The barcode and index sequences are listed in Table S10.

### RNA-seq and analysis

RNA-seq was performed as previously described^28^ with slight modifications. After RiboGreen quantification and quality control by Agilent BioAnalyzer, 100-500 ng of total RNA with RIN values of 9.4-10 underwent polyA selection and TruSeq library preparation according to instructions provided by Illumina (TruSeq Stranded mRNA LT Kit; RS-122-2102). Samples were amplified by 8 cycles of PCR, barcoded, and run on a NovaSeq 6000 platform in a PE100 run, using the NovaSeq 6000 S2 Reagent Kit (200 Cycles) (Illumina). The reads were aligned to hg38 with STAR_2.5.1b and parameters ‘--outFilterMultimapNmax 20 --alignSJoverhangMin 8 -- alignSJDBoverhangMin 1 --outFilter-MismatchNmax 999 --outFilterMismatchNoverReadLmax 0.04 --alignIntronMin 20 --alignIntronMax 1000000 --alignMatesGapMax 1000000.’ Transcripts were quantified with RSEM v1.2.23. Further downstream analysis was performed using DESeq2 (1.38.3).

